# DeepCellState: an autoencoder-based framework for predicting cell type-specific transcriptional states induced by drug treatment

**DOI:** 10.1101/2020.12.14.422792

**Authors:** Ramzan Umarov, Yu Li, Erik Arner

## Abstract

Drug treatment induces cell type-specific transcriptional programs, and as the number of combinations of drugs and cell types grows, the cost for exhaustive screens measuring the transcriptional drug response becomes intractable. We developed DeepCellState, a deep learning autoencoder-based framework, for predicting the induced transcriptional state in a cell type after drug treatment, based on the drug response in another cell type. Training the method on a large collection of transcriptional drug perturbation profiles, prediction accuracy improves significantly over baseline and alternative deep learning approaches when applying the method to two cell types, with improved accuracy when generalizing the framework to additional cell types. Treatments with drugs or whole drug families not seen during training are predicted with similar accuracy, and the same framework can be used for predicting the results from other interventions, such as gene knock-downs. Finally, analysis of the trained model shows that the internal representation is able to learn regulatory relationships between genes in a fully data-driven manner.

## Introduction

The transcriptional response to drug treatment is cell type-specific, with some drugs eliciting similar effects across lineages and others evoking a range of responses depending on the cell type^1,2^. High throughput profiling of the transcriptional effects of drug treatment has proven to be useful for analysis of drug mode of action^3^, drug repurposing^4^, and predicting off-target effects from drug treatment^5^. Although large repositories of gene expression profiles for a multitude of drug treatments in multiple cell types are available^6,7^, it is not combinatorially feasible to profile all the existing drugs in all the relevant cell types, motivating a need for methods that can accurately predict cell type-specific drug responses.

An autoencoder neural network is an unsupervised learning algorithm that finds an efficient compact representation of data by compressing and then reconstructing the original input. The primary goal of an autoencoder is dimensionality reduction. Autoencoders can have different architectures. However, the crucial feature is a bottleneck layer (latent layer), which has lower dimensionality compared to the input layer. Because of the bottleneck, only important features are captured by the model. Combining this property together with the addition of noise to the input allows for the construction of denoising autoencoders to build robust models from high-dimensional data. A number of recent studies have successfully applied autoencoders to biological problems, where deep autoencoders were used to denoise single-cell RNA-seq data sets^8,9^, analyze^10,11^ and predict^12,13^ gene expression, and to study the transcriptomic machinery^14^. Autoencoders have been applied to perturbation response modeling as well, focusing on single-cell data^15^, where for each perturbation, a large number of expression profiles are available with relatively low variance within sets of profiles from the same perturbation.

A particular application of autoencoders is DeepFake technology, mainly used in image and video processing applications to generate synthetic media where the likeness of one person is replaced by that of another one by training an autoencoder to compress the original input into a lower dimension latent space^16^. The same encoder part is used to compress media depicting two or more people, whereas separate decoders are used to decompress the likeness of each person (**Supplementary Figure 1**). This enables similar facial expressions or actions to be encoded in a similar way in the latent space while at the same time allowing for the reconstruction of person-specific facial details on the decoder side.

Considering that deep learning methods to date have not been widely applied to transcriptome perturbation analysis and prediction based on relatively small data sets with high variance, where each measurement represents a distinct state, we developed DeepCellState, an autoencoder-based framework inspired by DeepFake architecture, for predicting cell type-specific transcriptional drug response. Using a common decoder and separate decoders for each cell type (**Figure 1a**), the method accurately predicts the response in one cell type based on the response in another, with prediction accuracy increasing as the framework is generalized to multiple cell types. DeepCellState achieves significant improvement over the baseline and alternative deep learning approaches. Cell types not used in training can also be used as the basis for prediction, and the system is also able to predict the response of the entire drug families not seen by the network. Additionally, the same framework can be applied to predict the effects of loss of function (LoF) experiments. Analyzing the trained network, we find that it captures physical interactions between transcription factors and the target genes they regulate.

**Fig 1.**
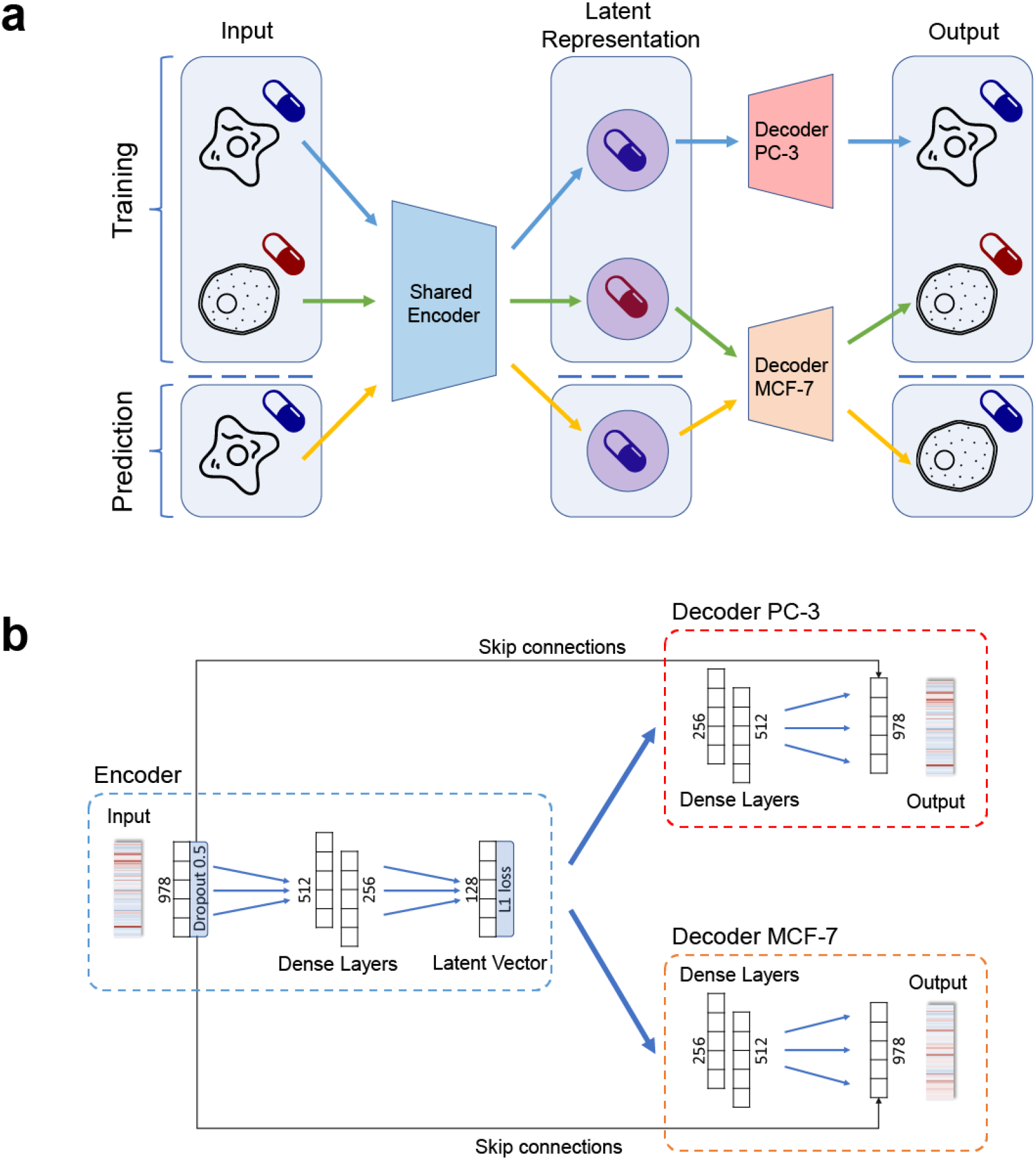
Overview of the proposed framework. **a**. Training and prediction procedures of DeepCellState. Transcriptional profiles are encoded by a shared encoder that captures the drug response in a cell neutral way. The latent representations of the drug responses are decoded by cell-specific decoders, which reconstruct the original input in a cell type-specific way. After the shared encoder and decoders are trained, the response to a drug in a cell type can be predicted by using the drug’s response in another cell type. **b**. The architecture of the encoder and the decoders used in DeepCellState. Dropout is applied to the input layer, forcing the model to denoise the input. The input is encoded as a vector of size 128 with a sparsity constraint enforced by L1 regularization. The model also employs skip connections from the dropout layer directly to the output layer, allowing some of the input to be directly copied to the output during training.

## Results

### DeepCellState network architecture for predicting cell type-specific drug response

The network architecture of DeepCellState, featuring a common encoder and cell type-specific decoders, is shown in **Figure 1b**. The encoder part uses two dense layers, with dropout applied to the input for increased generalization capabilities by forcing the model to denoise the input^17^. This is followed by a latent vector, which in turn is the input for the decoder, consisting of two layers with the same dimensions as the dense layers in the encoder. The encoder side is trained using transcriptional drug response data from multiple cell types, whereas distinct decoders are trained for each cell type separately. Additional details about the network architecture and training strategy are provided in **Methods**.

### Initial training and evaluation of model on two cell types

We initially developed the framework for two cell types, retrieving 12737 and 12031 transcriptional profiles from the LINCS database^6^ measuring the drug perturbation response in MCF-7 and PC-3 cell lines respectively, to use for initial training and evaluation (**Methods**). The transcriptional response profiles, measured at 978 “landmark genes” on the L1000 platform, were averaged across treatment time and doses in order to decrease noise^3^. We only kept profiles of drugs that were available for both MCF-7 and PC-3, resulting in 1750 averaged profiles for each cell type.

A naïve baseline method for predicting the transcriptional response in a cell type after treatment with a specific drug is to predict that the response to that drug will be the same as in another cell type. After training our network and using 10-fold cross-validation, holding out 10% of drugs, we obtained an average Pearson correlation coefficient (PCC) of 0.60 between predicted and actual treatment response to drugs not seen by the network, a significant improvement (p<1e-300, t-test) compared to the baseline average correlation of 0.28 between the responses in the input and output cell types (**Figure 2a,b,c,d, Supplementary Figure 2**), corresponding to an average fold-increase of 2.14 over baseline (**Figure 2b**). To make sure that the training was not biased by being exposed to drugs with similar targets, we next revised the testing strategy to hold out entire drug families with the same modes of action^18^, yielding similar results (**Supplementary Figure 3a, Supplementary Table 1, Methods**).

**Fig 2.**
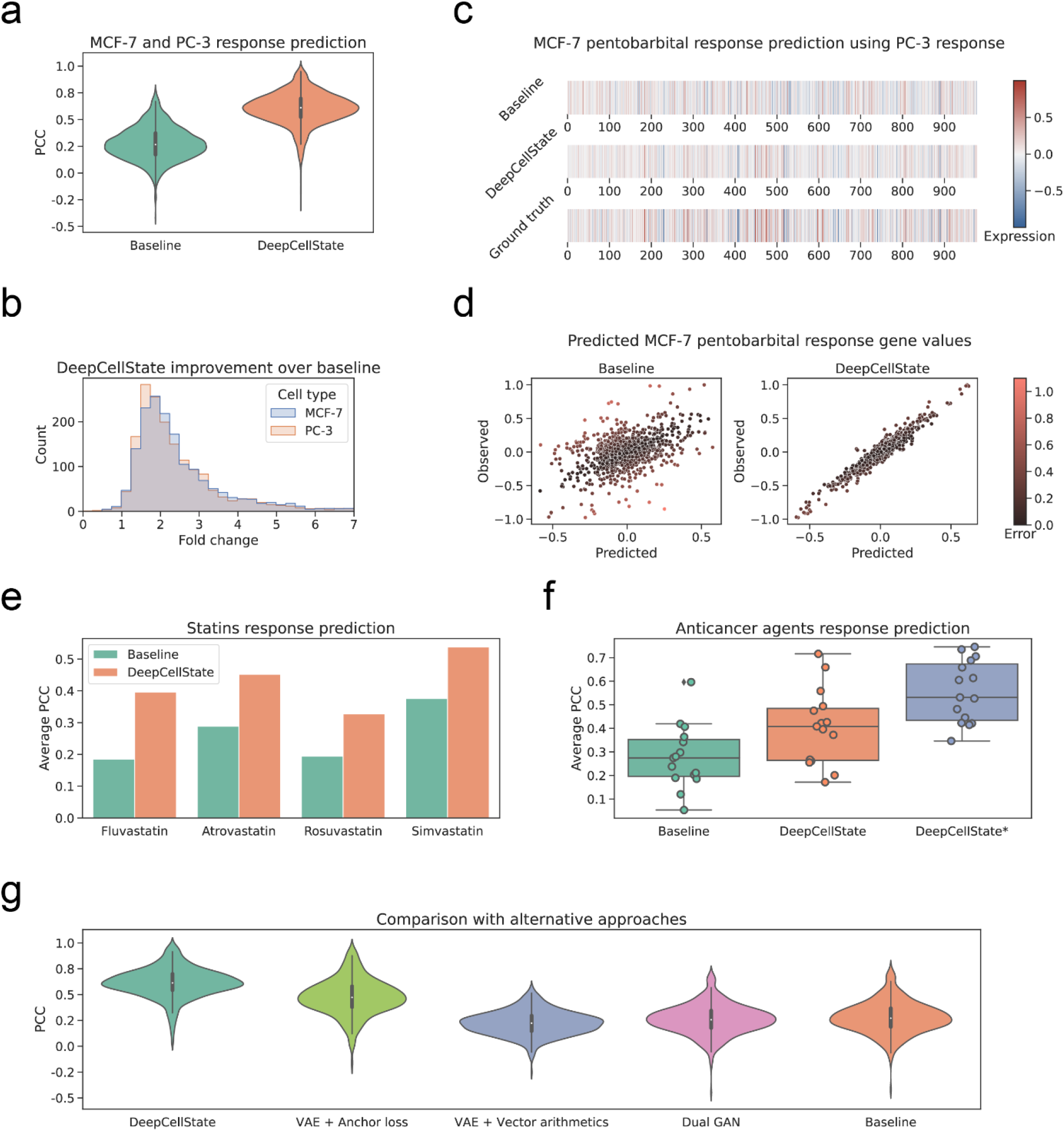
Evaluation of the DeepCellState performance. **a**. 10-fold cross-validation results. DeepCellState performance compared to the baseline. **b**. Distribution of fold-change in DeepCellState performance over baseline for predicting MCF-7 (blue) and PC-3 (red) response. **c**. From top to bottom pentobarbital response in PC-3 (baseline), MCF-7 pentobarbital response predicted by DeepCellState based on the PC-3 response, and the observed MCF-7 response (ground truth). **d**. Scatter plot of individual gene values predicted for pentobarbital response compared to the observed response in MCF-7 cells. **e**. Comparison of DeepCellState and baseline performance for prediction of statin response from an external data set. **f**. Comparison of predicted anticancer drug responses. DeepCellState* represents results obtained by applying transfer learning: performing additional training on 14 drugs and predicting one which is repeated 15 times each time leaving out one drug. **g**. Comparison with alternative methods for paired data modeling, using PC-3 responses to predict MCF-7 samples.

### Generalization of the method to multiple cell types

Proceeding to generalize the method to multiple cell types, we incrementally added transcription profiles from A375, HA1E, HT29, HELA, and YAPC cells (1796, 1796, 1750, 1570, and 1570 number of profiles respectively after averaging across treatment times and dosages), obtaining increased performance when the prediction was averaged over multiple input cell types compared to the two cell types case (PCC=0.66, p=1.88e-59, **Supplementary Figure 3b**). In comparison, a previously developed tensor-based method treating cell type-specific response prediction as an imputation problem^2^ was reported to achieve an average correlation of 0.54 using multiple cell types as input. We also tested the prediction capability using completely unseen cell types as input, obtaining lower performance, yet still significantly higher than baseline (**Supplementary Figure 3c**, PCC=0.49 compared to baseline PCC=0.28, p<1e-300).

### Evaluation of trained model on external data and application of transfer learning

To further validate the performance of DeepCellState, we evaluated its predictive capabilities on independently generated data sets on expression profiling platforms other than the L1000 platform used for the training data. In the first dataset we used, the expression profiles of four statins were measured in HepG2, MCF-7, and THP-1 cells^19^ using CAGE technology. Using HepG2 profiles as input and predicting MCF-7 response, the DeepCellState predictions showed a correlation of 0.43 with the true response in MCF-7 cells (1.64-fold over baseline, p=0.037, **Supplementary Table 2, Figure 2e**). Next, we tested our method using data from a study measuring the response to 15 anticancer agents^20^ on Affymetrix Human Genome U133A 2.0 Array platform, achieving 0.41 PCC (**Figure 2f)** compared to 0.28 PCC for baseline (1.46-fold improvement, p=0.026, **Supplemental Table 3**). Performance on this external dataset could be further improved through transfer learning, performing additional training of our model on a subset of the external data set. To this end, we used the model trained on the LINCS data and additionally trained it on 14 drugs from the anticancer set to predict the effects of treatment with the 15th drug. This was repeated 15 times, each time leaving out one drug, resulting in an average performance of 0.56 PCC (p=4.04e-06 compared to baseline). This result suggests that the transfer learning approach is useful to complete a small drug response data set which is not large enough to train the deep learning model by itself.

### Prediction of transcriptional effects of loss of function

We further hypothesized that the DeepCellState approach may be useful for predicting the response to cellular perturbations other than drug treatment. To test this, we added transcriptional profiles from loss of function (LoF) experiments from the same database to the training set. Using the updated set, we obtained an average correlation of 0.56 (corresponding to an average fold-increase over baseline of 1.6, p<1e-300, **Supplementary Figure 3d**) on the LoF experiments, the lower performance compared to drug perturbations likely due to that the number of available profiles for LoF was substantially lower than the number of drug treatment profiles (10230 LoF profiles in total compared to 24768 drug perturbation profiles for MCF-7 and PC-3 before averaging).

### Comparison with alternative approaches

We benchmarked DeepCellState against other potential methods to predict the cell response given the response in another cell. For this purpose, we focused on paired MCF-7 and PC-3 samples, with 30% data used for the test set. Recently, variational auto-encoders (VAEs) were used to deal with the paired biological data^21^. Anchor loss was used in this approach to ensure that paired samples are encoded closely in the latent space. Generative adversarial networks (GANs) were also applied to paired and semi-paired biological data^22,23^. The authors of MAGAN^23^ proposed a dual GAN setup with a special correspondence loss for the labeled data. In the dual GAN framework, each generator learns a mapping from one type of data to another in order to generate increasingly realistic output that cannot be distinguished from real data by a discriminator.

scGen^15^ was proposed to predict single-cell perturbation responses based on VAE architecture, where an encoder is used to learn the semantic embedding of both the normal cell state and the stimulated state. The difference between such embeddings could then be considered as the embedding of the drug stimulation. When applying the method onto new cell types, the authors proposed to perform latent space vector arithmetic, adding the stimulation embedding to the normal cell embedding, and to pass the resulting vector through the decoder, obtaining the predicted responses. Since we have multiple drugs and the amount of data for each drug is limited, we modified the method to learn the difference between different cell types (**Methods**).

Comparison of the tested methods on the benchmark dataset is shown in **Figure 2g**. We were unable to obtain a good performance using the dual GAN approach (PCC=0.26), likely due to the training data being too small. The data for each cell type has very high variance, making it hard for a discriminator to learn the difference between simulated and real cell response. scGen achieved an average correlation of 0.22, the main reason being that scGen is designed specifically for one kind of stimulation. However, in the problem we are trying to solve, multiple drugs and multiple cells need to be handled simultaneously and scGen showed limited capacity in handling this multi-task problem. The VAE using anchor loss obtained significant improvement over the baseline (PCC=0.48 compared to baseline PCC=0.28), however, the profiles predicted by DeepCellState were considerably more accurate (PCC=0.61, p<2.09e-40). After these experiments, we attempted to incorporate VAE as well as GAN into the DeepCellState approach, but the improvement was not significant enough to justify the added complexity.

### Effect of training set size and latent layer size on performance

We next investigated more deeply the relationship between the number of input profiles used in training and the prediction performance. Results showed that significant improvement over baseline was possible already for a small number of training examples, with higher performance as the number of training examples increased (**Figure 3a**).To study the impact of the latent layer size on the performance of DeepCellState, we tested five different layer sizes (8, 16, 32, 64 and 128 neurons) and trained our model 10 times for each size. Average results for the test set (shown in **Figure 3b**) showed that performance started to drop significantly at a latent layer size below 32.

**Fig 3.**
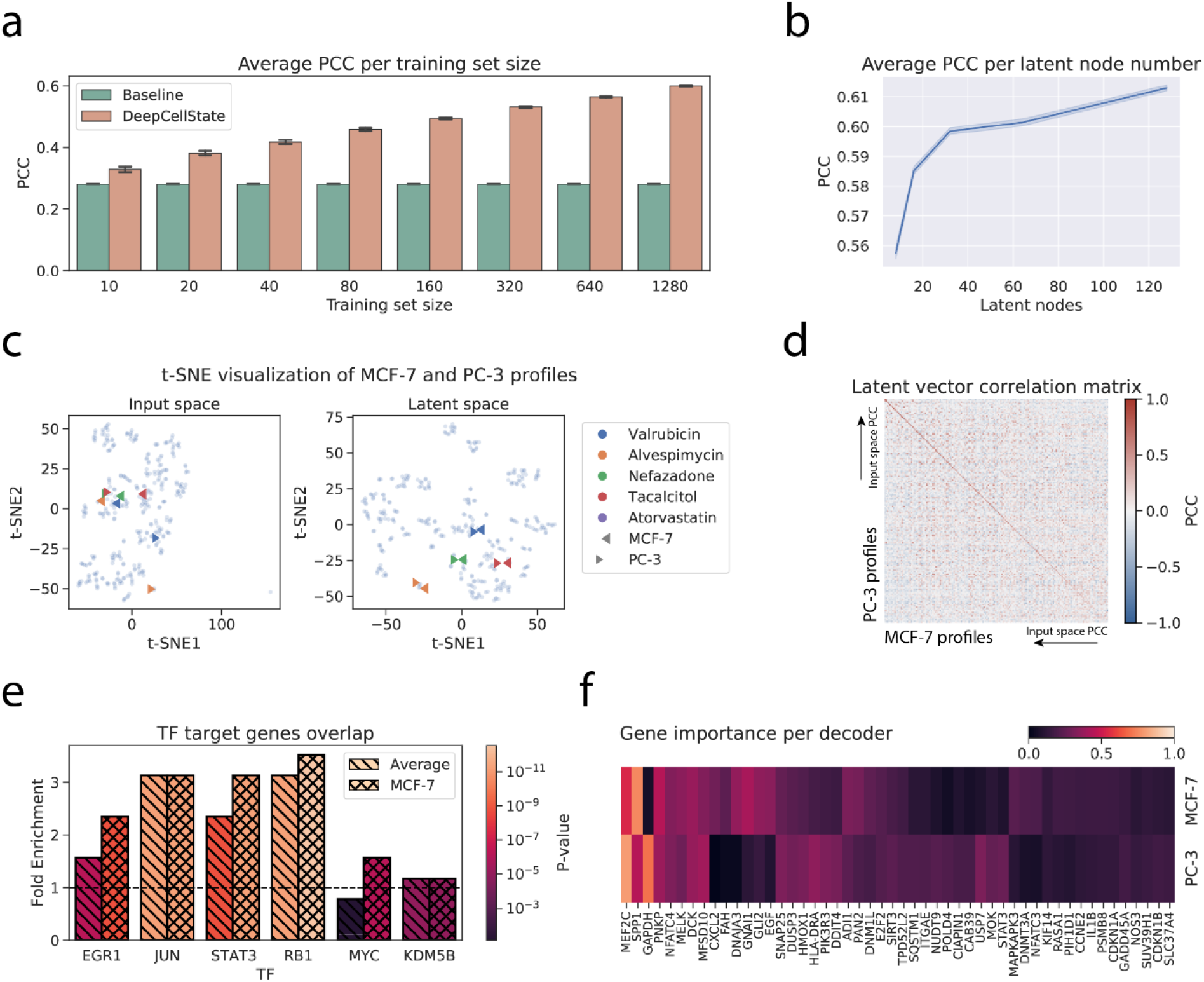
Analysis of the developed method. **a**. Average PCC obtained for different size subsets of the training set. **b**. Dependency between the number of latent nodes and the average PCC. **c**. t-SNE clustering of profiles in the input and the latent space. **d**. Matrix displaying latent vector response correlation. Drugs on each axis are sorted from high to low correlation in the input space. **e**. Overlaps of top target genes of TFs based on chromatin immunoprecipitation data compared to target gene prediction based on DeepCellState trained model. **f**. Heatmap of most important genes for MCF-7 and PC-3 decoders.

### Biological interpretation of trained model

We set out to analyze general and cell type-specific regulatory properties captured by the trained model, expecting that the latent vector itself should capture a general response to the input perturbations, whereas the decoder parts should represent behavior related to functionality specific to each cell type. To this end, we computed the PCC between the output of the nodes in the latent layer for the drugs in the test set between MCF-7 and PC-3 cells. Indeed, although considerably different, the inputs from the same drug in the two cell types consistently yielded similar output in the latent layer as illustrated in a t-SNE plot (**Figure 3c)** in the input and the latent space, where drugs with profiles divergent for different cell types in the input space (**Figure 3c**, left panel) could be observed to be encoded using very similar latent representations (**Figure 3c**, right panel). Similarly, a matrix containing correlations between latent responses for MCF-7 and PC-3 cells, sorted by the magnitude of the correlation of the input profiles, showed a consistently strong signal on the diagonal **(Figure 3d)**, independent of the correlation between the responses in the input space.

It has been shown previously that autoencoders are able to capture transcription factor (TF) gene regulatory relationships^14,24^. To test if this was the case also for the DeepCellState framework, we used TF target gene lists from ChIP-Atlas^25^ which include MCF-7 specific targets. After filtering, we computed the overlap between the targets implied by our model (see **Methods**) and targets based on ChIP-seq binding data, obtaining average fold enrichment of 2.5 (**Figure 3e**), with higher enrichment for the MCF-7 targets of four out of six transcription factors, compared to the general set of targets according to ChIP-Atlas. Identification of the most important genes in the cell type specific decoder response (see **Methods**) revealed enrichment of biological processes^26^ that matched known properties of each cell type (**Figure 3f**). For the top genes common to the decoders, the high ranking hits were enriched for terms including “response to abiotic stimulus” (p=0.0144) and “cellular response to chemical stimulus” (p=0.018), expected since the model was trained using drug profiles (**Supplemental Table 4**). Additionally, several significantly enriched terms (p=1.12e-04, p=2.32e-04) were related to positive regulation of cell death and apoptotic process, likely due to some doses being toxic as previously observed^27^. Conversely, top genes of the MCF-7 decoder included several genes involved in mammary gland development (SRC, STAT5B, EGF). Similarly, BMP4, GLI2, and FGFR2, which are known to be involved in prostate morphogenesis, were among the top important genes of the PC-3 decoder. Taken together, these results suggest that the trained network is able to capture known biological processes and regulatory relationships, and that additional, deeper analysis may reveal relationships previously unnoticed.

## Discussion

We here present DeepCellState, an autoencoder-based method that can successfully predict the transcriptional drug response in a cell type based on the response in another cell type. This problem was addressed in a recently proposed method that attempts to fill the gaps remaining in the combinatorial drug-cell space^2^ through a computational framework that first arranges existing profiles into a three-dimensional array (or tensor) indexed by drugs, genes, and cell types, and then uses either local or global information to predict unmeasured profiles. Different approaches were evaluated, with the best one achieving an average correlation of 0.54 compared to 0.66 for DeepCellState. One issue with the proposed tensor based method is that the performance strongly depends on the density of the constructed tensor, meaning that cell type-specific prediction of the response of a particular drug is heavily dependent on the amount of profiles for that same drug in other cell types. In contrast, the DeepCellState framework is capable of predicting the response of drugs never seen previously by the network and can also make predictions that improve significantly over baseline using the response in completely unseen cell types as input. Our results on even a relatively small amount of LoF data suggest that the approach may be able to capture changes in cellular state in a more general way.

Previous autoencoder-based methods for analyzing transcriptome and other data capturing cellular states have mainly focused on predicting cell state at the single-cell level^15,23,28^. Unlike these methods, DeepCellState can handle a large number of different perturbations, without requiring many data points for each. In a recent example, an autoencoder-based approach was used to translate between different domains of the single-cell data^21^, specifically between imaging and sequencing data. While the authors suggested the use of the approach for other types of translations, one requirement mentioned was that data come from the same cell population. With DeepCellState, we explicitly set out to predict the cell state in a particular cell type based on the state in one or more other cell types. Thus, we expect that the method can be readily applied to data sets measuring cell quantifiable characteristics other than transcriptome, e.g. imaging, Hi-C, or ATAC-seq data, with samples coming from different cell types.

The results here serve as a proof-of-principle and may be improved by various modifications to the architecture and training strategy. Additionally, the performance may be increased by measuring a larger set of genes than the 978 “landmark genes” measured on the L1000 platform, or by adding measurements of additional transcriptional features such as non-coding RNAs and transcribed enhancers. Performance is also expected to increase as more data capturing additional cellular states becomes available, especially if produced under controlled and standardized experimental conditions.

## Methods

### LINCS data processing

Drug response expression profiles from LINCS phase 2 dataset (GSE70138) were used for training and evaluation. The latest level 5 data (signatures from aggregating replicates and converted to z-scores) was used (downloaded May 27, 2020). There are usually several treatment times and doses for each drug treatment; we averaged the profiles for the same drug across different timepoints and doses to reduce noise. Finally, the profiles were normalized to the range from −1 to +1 based on the whole data matrix.

### Model architecture and training

The architecture of the autoencoder used in our method is shown in **Figure 1b**. The encoder consists of two dense layers with 512 and 256 neurons respectively. Dropout is applied to the input to increase the generalization capabilities of the model. The dropout rate is 0.5 which means half of the input genes will be set to 0 during training. This makes our model act as a denoising autoencoder since all values in the output layer need to be predicted. The last layer of the encoder is flattened and fed into a dense layer with 128 nodes, which represents our latent vector. The decoder takes the latent vector as an input and consists of two layers with 256 and 512 neurons respectively. The output layer has a direct connection to the dropout layer. The motivation is that some of the input can be “leaked” into the output^29^.

We use L1 regularization for the latent layer, which acts as a sparsity constraint on the activity of the latent representations. Activation function is leaky relu^30^ with alpha equal to 0.2 for all layers except the output layer, which uses tanh activation. The optimizer used is ADAM^31^ with a learning rate equal to 1e-4 and batch size is 128. Validation set is employed for early stopping with patience of 5 epochs.

To initialize our model, we first train a regular autoencoder using profiles of all cell types as both input and output. Then we create a separate decoder for each cell type by copying the weights from the autoencoder. These decoders are trained one by one together with the shared encoder, where the inputs are profiles from all cell types and the outputs are profiles with the decoder’s cell type only. In the final stage, when there has been no improvement in the validation set for 5 epochs, the encoder weights are frozen and only decoders are trained until there is no improvement. When making a prediction for a certain drug and cell combination, any cell profile can be used as an input. The cell specific decoder is used to make the final prediction.

DeepCellState is implemented in Python 3 using TensorFlow^32^ library and the models were trained using NVIDIA Quadro RTX 6000 GPU. Training two cell decoders takes about one hour to complete, however new predictions can be made in less than a second.

### External data processing

We trained an encoder and three decoders (HepG2, MCF-7 and PC-3) for external validation. The common drugs from the two external data sets were removed from the training and validation sets. The statin data set was generated by converting CAGE data to gene expression data by mapping to GENCODE version 34. CAGE tags in a 1000 base pair window around a landmark gene start were summed. We added a small value c = 2 to each gene value to avoid problems when calculating log fold change. Each generated profile was multiplied by a million and divided by the total sum of the profile. The resulting profiles for controls and treatments were averaged across replicates followed by a computation of log fold change after treatment for each gene. For the anticancer data we directly calculated log fold change between the average control and treatment values. In both statin and anticancer tests, we only used landmark genes, setting missing gene values to 0. Only the 24 h time point was used as it is the most common time point in LINCS phase 2 compound data (96% of the data).

### Methods related to analysis/interpretation of model

Since it is hard to directly learn the inner-workings of the deep learning model, we study it by feeding in different inputs and analyzing the output. To calculate the top genes for each decoder, we iterated through all the data, for each profile using 100 random genes subsets of size 100 to input in the model. The gene sets which performed the best to make the prediction were considered to be important. For each profile we picked the top 10 subsets that performed the best. After doing this for all the profiles, we ranked the genes based on their frequency in these subsets.

For the TF analysis, we downloaded target genes for 22 TFs profiled in MCF-7 cells from ChIP-Atlas^25^. We filtered them by retaining TFs that were upregulated in at least one drug treatment profile (i.e. has value > 0.5), and that had at least 50 targets in ChIP-Atlas (with targets defined as genes having a binding score of at least 100). This resulted in 6 remaining TFs and we picked the top 50 target genes for each. To make a prediction of target genes using DeepCellState, we started with a random profile and performed gradient descent optimization to find the input profile that maximizes the TF. As the obtained profile will also have other upregulated genes, the top 50 with the highest values are the DeepCellState prediction for the TF targets. For each TF we calculated fold enrichment and p-values using binomial statistic for the overlap between the DeepCellState set and the ChIP-Atlas set, using a similar method as described on the PANTHER website^26^.

### Comparison with alternative approaches

scGen^15^ was designed to predict single-cell perturbation responses based on VAE architecture, where an encoder *f* is used to learn the semantic embedding, z, of both the normal cell state and the stimulated state. The difference between such embeddings could then be considered as the embedding of the drug stimulation. When applying the method onto new cell types, the authors proposed to perform latent space vector arithmetic, adding the stimulation embedding, δ, to the normal cell embedding, and to pass the resulting vector through the decoder, *h*, obtaining the predicted responses.

Mathematically, we have

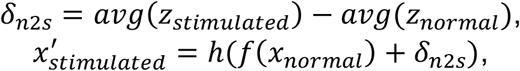

where δ_*n*2*s*_ is the stimulation embedding; *z*_*stimulated*_ is the embedding of the stimulated cell profile; *z*_*normal*_ is the embedding of the normal cell profile; *x*_*normal*_ is the input normal cell profile; 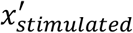 is the predicted stimulated cell profile.

Regarding our task, since we have multiple drugs and the amount of data for each drug is limited, we modified the method to learn the difference between different cell types, instead of the difference between conditions.

Consequently, we have

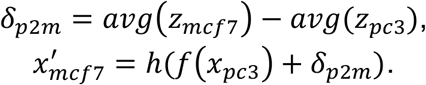

Both the encoder, *f*, and decoder, *h*, are trained using the standard VAE training protocol described in the original paper^15^ with our data as discussed above.

MAGAN^23^ was developed to tackle the problem of aligning corresponding sets of samples. MAGAN is composed of two GANs, each with a generator network *G* that takes as input *X*and outputs a target dataset *X*^′^. For the minibatch (*x*_1_,*x*_2_), the loss for generator *G*_12_ is defined as sum of reconstruction loss, discriminator loss, and correspondence loss:

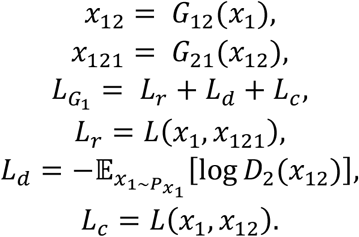

The loss for discriminator *D*_1_ is defined as:

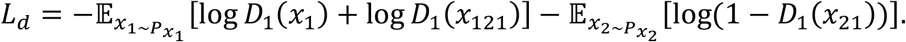

Since in our task all the data are paired, we used the following correspondence loss, as suggested in the original paper^23^:

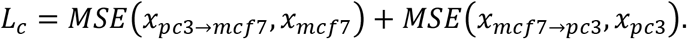

We used the total loss to train two generators, *G*_*pc*3→*mcf*7_and *G*_*mcf*7→*pc*3_, together with two discriminators, *D*_*pc*3_ and *D*_*mcf*7_.

To implement VAE with anchor loss, we have trained a standard VAE with two decoders and employed the following loss to ensure that all the *m* paired samples are encoded close to each other by encoder *E*in the latent space:

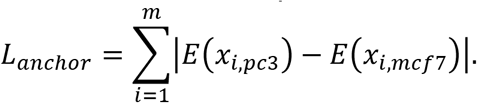

## Data and code availability

Trained models, code to generate them, and code for analysis and figures described in this study are available at the following GitHub repository: https://github.com/umarov90/DeepCellState. All data used in training and validation is publicly available through databases referenced above.

## Acknowledgements

We would like to thank Andrew Tae-Jun Kwon and Bogumil Kaczkowski for insightful comments on the manuscript.

## Author contributions

EA conceived the study and supervised the research. RU performed the research and drafted the manuscript together with EA. YL performed comparisons with alternative approaches.

