## Supplemental Information for "DeepCellState: an autoencoder-based framework for predicting cell type-specific transcriptional states induced by drug treatment"

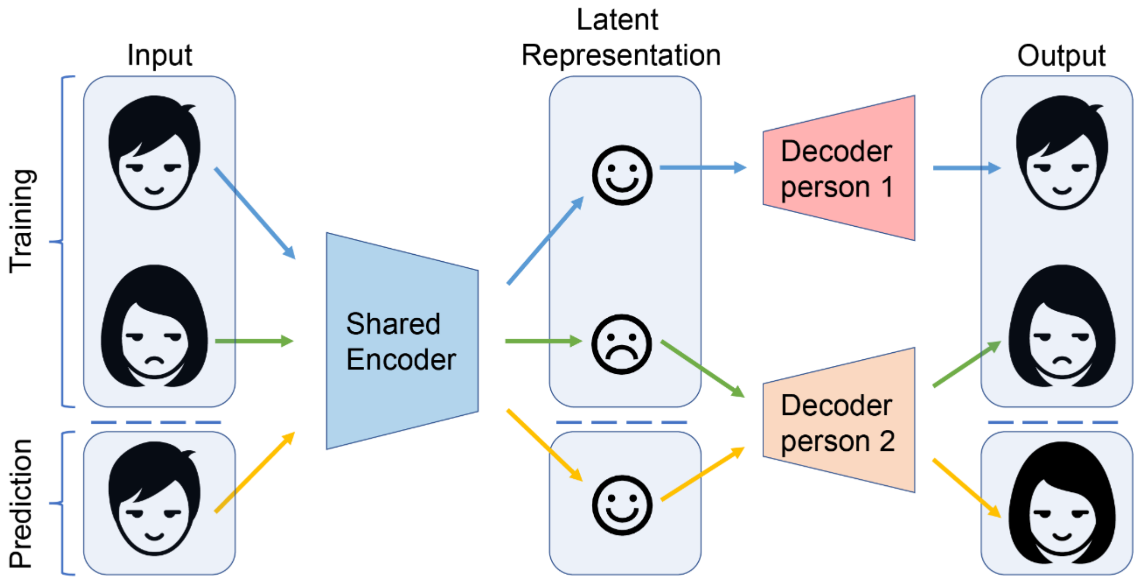


**Fig. S1** DeepFake approach for the face swap. Similar facial expressions are encoded in a similar way in the latent space, while person specific facial details are reconstructed on the decoder side.

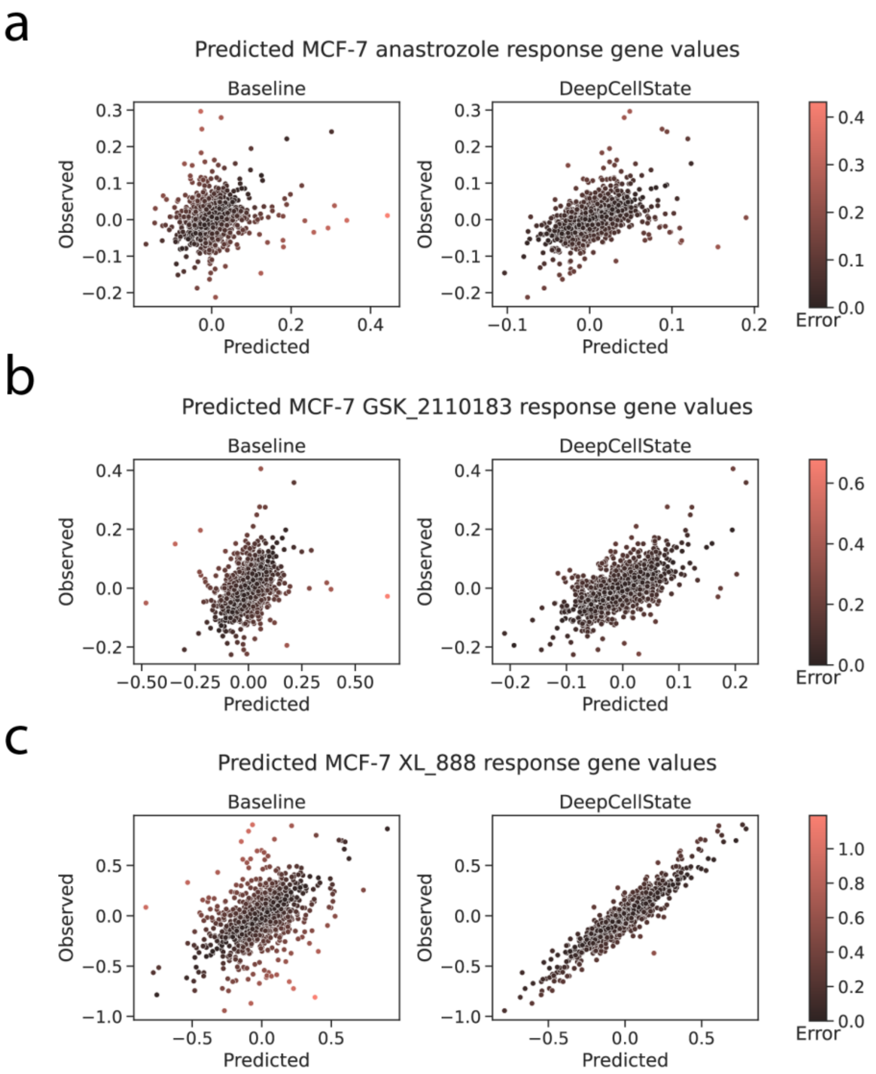


**Fig. S2** Examples of MCF-7 profiles predicted with low, medium and high PCC. **a.** MCF-7 anastrozole response predicted with PCC 0.47. **b.** MCF-7 GSK 2110183 response predicted with 0.61 PCC. **c.** MCF-7 XL 888 response predicted with PCC 0.93.

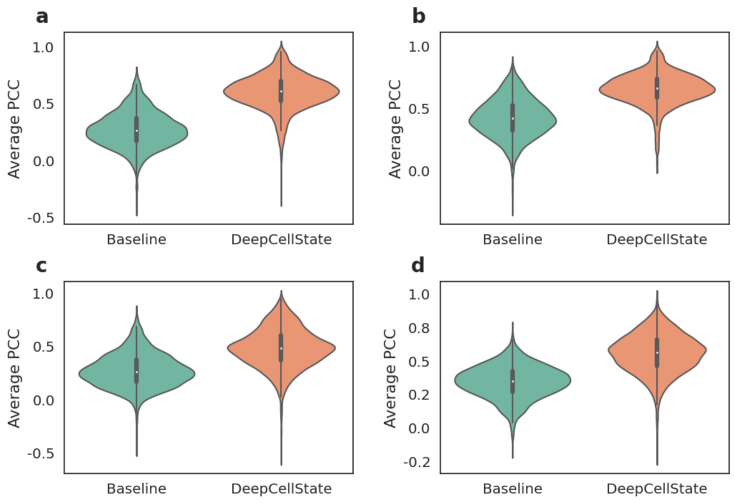


**Fig. S3** Different testing strategies of DeepCellState. **a** Results obtained by holding out entire drug families for the testing. **b** Performance of DeepCellSate by inclusion of profiles from other cell types in the training set. **c** Performance evaluation using completely unseen cell type as input. **d** Results for shRNA for LoF experiments profiles prediction.

| Family | Count |
|---|---|
| antibiotics | 161 |
| adrenergic | 112 |
| cholinergic | 101 |
| 5-HT modulator | 89 |
| TKI | 74 |
| COX inh. | 73 |
| histaminergic | 68 |
| antipsychotic | 55 |
| GABAergic | 54 |
| dopaminergic | 51 |

**Table S1** Number of drugs in the ten drug families with the most number of drugs.

| Drug | Baseline | DeepCellState |
| --- | --- | --- |
| Fluvastatin | 0.19 | 0.40 |
| Atrovastatin | 0.29 | 0.45 |
| Rosuvastatin | 0.19 | 0.33 |
| Simvastatin | 0.38 | 0.54 |
| Total | 0.26 | 0.43 |

**Table S2** Performance of Baseline and DeepCellState when predicting statin response profiles.

| Drug | Baseline | DeepCellState | DeepCellState* |
| --- | --- | --- | --- |
| 5-Azacytidine | 0.20 | 0.17 | 0.60 |
| bortezomib | 0.42 | 0.66 | 0.70 |
| cisplatin | 0.30 | 0.42 | 0.74 |
| dasatinib | 0.12 | 0.26 | 0.53 |
| doxorubicin | 0.19 | 0.47 | 0.64 |
| erlotinib | 0.27 | 0.37 | 0.74 |
| geldanamycin | 0.28 | 0.56 | 0.60 |
| gemcitibine | 0.24 | 0.27 | 0.41 |
| lapatinib | 0.05 | 0.20 | 0.37 |
| paclitaxel | 0.41 | 0.41 | 0.41 |
| sirolimus | 0.34 | 0.49 | 0.50 |
| sorafenib | 0.36 | 0.43 | 0.53 |
| sunitinib | 0.21 | 0.25 | 0.46 |
| topotecan | 0.19 | 0.40 | 0.43 |
| vorinostat | 0.60 | 0.72 | 0.69 |
| Total | 0.28 | 0.41 | 0.56 |

**Table S3** Performance of Baseline and DeepCellState when predicting anticancer agents response profiles. DeepCellState* are results obtained after performing additional training on 14 anticancer agents and predicting for one repeated for each drug.

| GO biological process complete | # | # | expected | Fold Enrichment | +/- | raw P value |
|---|---|---|---|---|---|---|
| programmed cell death | 130 | 18 | 7.07 | 2.55 | + | 1.12E-04 |
| cell death | 131 | 18 | 7.12 | 2.53 | + | 1.24E-04 |
| apoptotic process | 125 | 17 | 6.79 | 2.50 | + | 2.32E-04 |
| muscle adaptation | 6 | 4 | .33 | 12.27 | + | 3.31E-04 |
| intracellular signal transduction | 202 | 22 | 10.98 | 2.00 | + | 5.14E-04 |
| smooth muscle adaptation | 3 | 3 | .16 | 18.40 | + | 6.09E-04 |
| negative regulation of smooth muscle cell proliferation | 13 | 5 | .71 | 7.08 | + | 7.13E-04 |
| negative regulation of vascular associated smooth muscle cell proliferation | 8 | 4 | .43 | 9.20 | + | 9.66E-04 |
| mitotic cell cycle arrest | 4 | 3 | .22 | 13.80 | + | 1.39E-03 |
| response to lipid | 107 | 14 | 5.82 | 2.41 | + | 1.41E-03 |
| positive regulation of cell death | 99 | 13 | 5.38 | 2.42 | + | 2.10E-03 |
| regulation of macromolecule metabolic process | 497 | 38 | 27.01 | 1.41 | + | 2.15E-03 |
| regulation of cell death | 224 | 22 | 12.17 | 1.81 | + | 2.17E-03 |

| | | | | | | |
|---|---|---|---|---|---|---|
| negative regulation of biological process | 461 | 36 | 25.05 | 1.44 | + | 2.33E-03 |
| intrinsic apoptotic signaling pathway | 34 | 7 | 1.85 | 3.79 | + | 2.38E-03 |
| cellular response to organic cyclic compound | 65 | 10 | 3.53 | 2.83 | + | 2.42E-03 |
| cellular response to stimulus | 519 | 39 | 28.21 | 1.38 | + | 2.44E-03 |
| defense response | 101 | 13 | 5.49 | 2.37 | + | 2.51E-03 |
| apoptotic signaling pathway | 55 | 9 | 2.99 | 3.01 | + | 2.70E-03 |
| inflammatory response | 48 | 8 | 2.61 | 3.07 | + | 4.24E-03 |
| neutrophil migration | 6 | 3 | .33 | 9.20 | + | 4.34E-03 |
| granulocyte chemotaxis | 6 | 3 | .33 | 9.20 | + | 4.34E-03 |
| neutrophil chemotaxis | 6 | 3 | .33 | 9.20 | + | 4.34E-03 |
| granulocyte migration | 6 | 3 | .33 | 9.20 | + | 4.34E-03 |
| modulation of chemical synaptic transmission | 38 | 7 | 2.07 | 3.39 | + | 4.40E-03 |
| regulation of trans-synaptic signaling | 38 | 7 | 2.07 | 3.39 | + | 4.40E-03 |
| regulation of anatomical structure morphogenesis | 108 | 13 | 5.87 | 2.21 | + | 4.50E-03 |
| negative regulation of apoptotic process | 137 | 15 | 7.45 | 2.01 | + | 5.31E-03 |

| | | | | | | |
|---|---|---|---|---|---|---|
| calcineurin-mediated signaling | 2 | 2 | .11 | 18.40 | + | 5.41E-03 |
| arginine catabolic process | 2 | 2 | .11 | 18.40 | + | 5.41E-03 |
| arginine metabolic process | 2 | 2 | .11 | 18.40 | + | 5.41E-03 |
| calcineurin-NFAT signaling cascade | 2 | 2 | .11 | 18.40 | + | 5.41E-03 |
| glutamine family amino acid catabolic process | 2 | 2 | .11 | 18.40 | + | 5.41E-03 |
| negative regulation of cardiac muscle tissue regeneration | 2 | 2 | .11 | 18.40 | + | 5.41E-03 |
| regulation of cardiac muscle tissue regeneration | 2 | 2 | .11 | 18.40 | + | 5.41E-03 |
| muscle hyperplasia | 2 | 2 | .11 | 18.40 | + | 5.41E-03 |
| smooth muscle hyperplasia | 2 | 2 | .11 | 18.40 | + | 5.41E-03 |
| positive regulation of cerebellar granule cell precursor proliferation | 2 | 2 | .11 | 18.40 | + | 5.41E-03 |
| regulation of cerebellar granule cell precursor proliferation | 2 | 2 | .11 | 18.40 | + | 5.41E-03 |
| negative regulation of glycolytic process | 2 | 2 | .11 | 18.40 | + | 5.41E-03 |
| negative regulation of macromolecule metabolic process | 273 | 24 | 14.84 | 1.62 | + | 5.70E-03 |
| regulation of secretion | 62 | 9 | 3.37 | 2.67 | + | 5.93E-03 |
| negative regulation of programmed cell death | 139 | 15 | 7.55 | 1.99 | + | 6.08E-03 |

| | | | | | | |
|---|---|---|---|---|---|---|
| negative regulation of developmental process | 99 | 12 | 5.38 | 2.23 | + | 6.13E-03 |
| negative regulation of cell death | 155 | 16 | 8.42 | 1.90 | + | 6.90E-03 |
| regulation of molecular function | 310 | 26 | 16.85 | 1.54 | + | 7.01E-03 |
| positive regulation of catalytic activity | 185 | 18 | 10.05 | 1.79 | + | 7.30E-03 |
| regulation of apoptotic process | 200 | 19 | 10.87 | 1.75 | + | 7.33E-03 |
| regulation of metabolic process | 545 | 39 | 29.62 | 1.32 | + | 7.38E-03 |
| biological regulation | 782 | 50 | 42.50 | 1.18 | + | 7.65E-03 |
| regulation of catalytic activity | 263 | 23 | 14.29 | 1.61 | + | 7.66E-03 |
| cell surface receptor signaling pathway | 232 | 21 | 12.61 | 1.67 | + | 7.88E-03 |
| regulation of multicellular organismal development | 143 | 15 | 7.77 | 1.93 | + | 7.90E-03 |
| negative regulation of cellular metabolic process | 264 | 23 | 14.35 | 1.60 | + | 8.04E-03 |
| negative regulation of metabolic process | 297 | 25 | 16.14 | 1.55 | + | 8.24E-03 |
| response to stress | 348 | 28 | 18.91 | 1.48 | + | 8.51E-03 |
| negative regulation of cellular process | 436 | 33 | 23.70 | 1.39 | + | 8.54E-03 |
| osteoblast differentiation | 15 | 4 | .82 | 4.91 | + | 9.03E-03 |

| | | | | | | |
|---|---|---|---|---|---|---|
| positive regulation of cellular process | 493 | 36 | 26.79 | 1.34 | + | 9.14E-03 |
| cellular response to lipid | 67 | 9 | 3.64 | 2.47 | + | 9.66E-03 |
| regulation of phosphate metabolic process | 206 | 19 | 11.20 | 1.70 | + | 1.01E-02 |
| regulation of programmed cell death | 206 | 19 | 11.20 | 1.70 | + | 1.01E-02 |
| regulation of phosphorus metabolic process | 206 | 19 | 11.20 | 1.70 | + | 1.01E-02 |
| cellular response to DNA damage stimulus | 93 | 11 | 5.05 | 2.18 | + | 1.05E-02 |
| negative regulation of nitrogen compound metabolic process | 238 | 21 | 12.93 | 1.62 | + | 1.06E-02 |
| response to organic cyclic compound | 107 | 12 | 5.82 | 2.06 | + | 1.12E-02 |
| regulation of endothelial cell proliferation | 16 | 4 | .87 | 4.60 | + | 1.12E-02 |
| response to heat | 16 | 4 | .87 | 4.60 | + | 1.12E-02 |
| regulation of vascular associated smooth muscle cell proliferation | 16 | 4 | .87 | 4.60 | + | 1.12E-02 |
| negative regulation of cellular protein metabolic process | 135 | 14 | 7.34 | 1.91 | + | 1.16E-02 |
| positive regulation of nucleotide biosynthetic process | 3 | 2 | .16 | 12.27 | + | 1.18E-02 |
| inositol phosphate-mediated signaling | 3 | 2 | .16 | 12.27 | + | 1.18E-02 |
| negative regulation of purine nucleotide metabolic process | 3 | 2 | .16 | 12.27 | + | 1.18E-02 |

| | | | | | | |
|---|---|---|---|---|---|---|
| positive regulation of purine nucleotide biosynthetic process | 3 | 2 | .16 | 12.27 | + | 1.18E-02 |
| negative regulation of cellular carbohydrate metabolic process | 3 | 2 | .16 | 12.27 | + | 1.18E-02 |
| negative regulation of nucleotide metabolic process | 3 | 2 | .16 | 12.27 | + | 1.18E-02 |
| acute-phase response | 3 | 2 | .16 | 12.27 | + | 1.18E-02 |
| cerebellar cortex morphogenesis | 3 | 2 | .16 | 12.27 | + | 1.18E-02 |
| cellular response to chemical stimulus | 305 | 25 | 16.58 | 1.51 | + | 1.18E-02 |
| regulation of protein phosphorylation | 179 | 17 | 9.73 | 1.75 | + | 1.19E-02 |
| negative regulation of protein modification process | 95 | 11 | 5.16 | 2.13 | + | 1.22E-02 |
| myeloid leukocyte migration | 9 | 3 | .49 | 6.13 | + | 1.31E-02 |
| negative regulation of catabolic process | 36 | 6 | 1.96 | 3.07 | + | 1.32E-02 |
| regulation of angiogenesis | 36 | 6 | 1.96 | 3.07 | + | 1.32E-02 |
| response to stimulus | 600 | 41 | 32.61 | 1.26 | + | 1.34E-02 |
| positive regulation of biological process | 522 | 37 | 28.37 | 1.30 | + | 1.35E-02 |
| cell projection organization | 110 | 1 | 5.98 | .17 | - | 1.38E-02 |

| | | | | | | |
|---|---|---|---|---|---|---|
| intrinsic apoptotic signaling pathway in response to DNA damage | 17 | 4 | .92 | 4.33 | + | 1.38E-02 |
| response to abiotic stimulus | 153 | 15 | 8.32 | 1.80 | + | 1.44E-02 |
| negative regulation of gene expression | 183 | 17 | 9.95 | 1.71 | + | 1.47E-02 |
| negative regulation of protein metabolic process | 139 | 14 | 7.55 | 1.85 | + | 1.47E-02 |
| signaling | 414 | 31 | 22.50 | 1.38 | + | 1.48E-02 |
| negative regulation of cellular component movement | 37 | 6 | 2.01 | 2.98 | + | 1.50E-02 |
| regulation of vasculature development | 37 | 6 | 2.01 | 2.98 | + | 1.50E-02 |
| negative regulation of multicellular organismal process | 98 | 11 | 5.33 | 2.07 | + | 1.52E-02 |
| regulation of secretion by cell | 60 | 8 | 3.26 | 2.45 | + | 1.53E-02 |
| signal transduction by p53 class mediator | 27 | 5 | 1.47 | 3.41 | + | 1.56E-02 |
| negative regulation of cellular macromolecule biosynthetic process | 140 | 14 | 7.61 | 1.84 | + | 1.56E-02 |
| signal transduction | 398 | 30 | 21.63 | 1.39 | + | 1.58E-02 |
| plasma membrane bounded cell projection organization | 107 | 1 | 5.82 | .17 | - | 1.61E-02 |
| signal transduction by protein phosphorylation | 61 | 8 | 3.32 | 2.41 | + | 1.68E-02 |

| | | | | | | |
|---|---|---|---|---|---|---|
| regulation of gene expression | 382 | 29 | 20.76 | 1.40 | + | 1.68E-02 |
| positive regulation of endothelial cell proliferation | 10 | 3 | .54 | 5.52 | + | 1.73E-02 |
| leukocyte chemotaxis | 10 | 3 | .54 | 5.52 | + | 1.73E-02 |
| regulation of peptidyl-threonine phosphorylation | 10 | 3 | .54 | 5.52 | + | 1.73E-02 |
| intrinsic apoptotic signaling pathway by p53 class mediator | 10 | 3 | .54 | 5.52 | + | 1.73E-02 |
| regulation of biological process | 755 | 48 | 41.03 | 1.17 | + | 1.75E-02 |
| positive regulation of phosphorus metabolic process | 142 | 14 | 7.72 | 1.81 | + | 1.76E-02 |
| positive regulation of phosphate metabolic process | 142 | 14 | 7.72 | 1.81 | + | 1.76E-02 |
| positive regulation of programmed cell death | 87 | 10 | 4.73 | 2.11 | + | 1.78E-02 |
| circulatory system development | 87 | 10 | 4.73 | 2.11 | + | 1.78E-02 |
| cell communication | 420 | 31 | 22.83 | 1.36 | + | 1.85E-02 |
| developmental process | 420 | 31 | 22.83 | 1.36 | + | 1.85E-02 |
| metabolic process | 670 | 44 | 36.41 | 1.21 | + | 1.85E-02 |
| negative regulation of macromolecule biosynthetic process | 143 | 14 | 7.77 | 1.80 | + | 1.86E-02 |
| negative regulation of molecular function | 129 | 13 | 7.01 | 1.85 | + | 1.89E-02 |

| | | | | | | |
|---|---|---|---|---|---|---|
| phosphorylation | 158 | 15 | 8.59 | 1.75 | + | 1.90E-02 |
| regulation of cellular component movement | 115 | 12 | 6.25 | 1.92 | + | 1.90E-02 |
| regulation of synaptic plasticity | 19 | 4 | 1.03 | 3.87 | + | 1.98E-02 |
| cytokine production | 19 | 4 | 1.03 | 3.87 | + | 1.98E-02 |
| positive regulation of small molecule metabolic process | 19 | 4 | 1.03 | 3.87 | + | 1.98E-02 |
| positive regulation of synaptic transmission | 19 | 4 | 1.03 | 3.87 | + | 1.98E-02 |

**Table S4**. Enriched Gene Ontology terms for top important genes in the latent layer.
